# Evoked resonant neural activity outperforms spectral markers in decoding sleep from the subthalamic nucleus

**DOI:** 10.1101/2024.08.20.608562

**Authors:** Christoph Wiest, Thomas G. Simpson, Alek Pogosyan, Harutomo Hasegawa, Shenghong He, Fernando Rodriguez Plazas, Laura Wehmeyer, Sahar Yassine, Xuanjun Guo, Rahul Shah, Anca Merla, Andrea Perera, Ahmed Raslan, Andrew O’Keeffe, Michael G. Hart, Francesca Morgante, Erlick A. Pereira, Keyoumars Ashkan, Huiling Tan

## Abstract

**Background:** Deep brain stimulation is a treatment for advanced Parkinson’s disease and currently tuned to target motor symptoms during daytime. Parkinson’s disease is associated with multiple nocturnal symptoms such as akinesia, insomnia and sleep fragmentation which may require adjustments of stimulation during sleep for best treatment outcome.

**Objectives:** There is a need for a robust biomarker to guide stimulation titration across sleep stages. This study aimed to investigate whether evoked resonant neural activity (ERNA) is modulated during sleep.

**Methods:** We recorded local field potentials from the subthalamic nucleus of four Parkinson’s patients with externalised electrodes while applying single stimulation pulses to investigate the effect of sleep on ERNA.

**Results:** We found that ERNA features change with wakefulness and sleep stages, and are correlated with canonical frequency bands and heart rate. We further evaluated the performance of machine learning models in classifying non-REM sleep versus wakefulness and found that ERNA amplitude outperforms all spectral markers.

**Conclusions:** Given the heterogeneity of spectral features during sleep, their susceptibility to movement artefacts and superior classification accuracy of models using ERNA features, this study paves the way for ERNA as a marker for automatic stimulation titration during sleep and improved patient care.

## 1. Introduction

Parkinson’s disease (PD) is the second most common neurodegenerative disorder and anticipated to double in prevalence over the next 30 years^1^. Apart from daytime motor symptoms, nocturnal distress and sleep impairment such as nocturnal akinesia, fragmented sleep, insomnia and decreased time in deep sleep and REM stages are a hallmark of PD^2^. High-frequency deep brain stimulation (DBS) of the subthalamic nucleus (STN) is an effective therapy for PD, but is titrated around daytime motor fluctuations and currently not optimised for night time or within sleep stages. Current closed-loop DBS algorithms are primarily using beta activity as a feedback signal which may lead to suboptimal therapy and sleep disruption as beta oscillation is naturally reduced at night relative to day^3^. Therefore, adaptive DBS that modulates stimulation parameters in response to detected sleep stages (known as sleep-aware adaptive DBS) may control nocturnal PD symptoms better. Automatic sleep stage classification based on cortical^4–6^ or subthalamic^7,8^ signals using different machine learning algorithms has been implemented in offline analysis with varying accuracy between ∼70% and ∼95%. Alternatively, a simpler approach is to adjust stimulation based on the time of day^9^, which neglects day-to-day variations in the time patients will go to bed or fall asleep. Here, we report for the first time that a neurophysiological marker in subthalamic local field potentials (LFP), i.e. stimulation-evoked resonant neural activity (ERNA), tracks sleep onset and sleep stage transitions, which may enhance and simplify automatic DBS titration at night.

## 2. Methods

### 2.1. Consent, regulatory approval and patient selection

This protocol was approved by the Health Research Authority UK and the local Research Ethics Committee (IRAS: 46576). Four patients with idiopathic PD undergoing bilateral STN-DBS surgery were recruited for LFP recording at King’s College Hospital NHS Foundation Trust, London, or St. George’s University Hospital NHS Foundation Trust, London. Written informed consent was obtained in line with the Declaration of the Principles of Helsinki. Patients were selected by an interdisciplinary team as described before^10^. The average age at recording was 63 ± 3.34 years (mean ± SEM) with average disease duration of 17.25 ± 4.31 years. Clinical details are summarised in **Table 1**.

**Table 1.** Patient information, recording details and classifier performance. UPDRS: Unified Parkinson’s Disease Rating Scale; DBS: deep brain stimulation; STN: subthalamic nucleus; R: right; L: left; SG: St. George’s Hospital; K: King’s College Hospital; Medt: Medtronic; Boston: Boston Scientific; WS: window size; AUC: area under the curve of the receiver operating characteristic; LR: logistic regression; RF: random forest; SVC: support vector machine; PRKN mut: heterozygous PRKN mutation.

| Patient #<br>(Hospital) | Gender<br>(m/f) | Age<br>(yr) | Disease<br>Duration<br>(yr) | UPDRS-III<br>OFF meds<br>(pre-OP) | UPDRS-III<br>ON meds<br>(pre-OP) | Predominant<br>Symptoms | Time of<br>Recording<br>(days post-OP) | DBS<br>system | STN<br>tested<br>(R/L) | Duration<br>sleep<br>recording | Best ERNA Classification<br>Accuracy (AUC) | Best Spectral<br>Classification Accuracy<br>(AUC) | Best Overall Classification<br>Accuracy (AUC) |
| --- | --- | --- | --- | --- | --- | --- | --- | --- | --- | --- | --- | --- | --- |
| 1 (SG) | m | 69 | 30 | 37 | 31 | tremor<br>(PRKN mut) | 4 | Boston dir | R | 1 h 31 min 56 sec | WS1: 87.35 (93.10) SVC<br>WS10: 95.35 (98.23) LR | WS1: 83.88 (85.43) LR<br>WS10: 84.98 (86.07) LR | WS1: 93.22 (98.04) LR<br>WS10: 96.94 (99.05) LR |
| 2 (K) | f | 67 | 12 | 57 | 9 | tremor | 4 | Medt SenSight | L | 32 min 58 sec | N/A | N/A | N/A |
| 3 (K) | m | 62 | 12 | 53 | 12 | tremor | 6 | Medt SenSight | L | 9 min 51 sec | 97.69 (99.76) LR | 90.19 (96.31) RF | 99.23 (100) LR |
| 4 (K) | m | 54 | 15 | 62 | 23 | tremor | 6 | Medt SenSight | R | 10 min 0 sec | 90 (100) SVC | 90 (100) SVC | 90 (100) LR |

### 2.2. Surgery and lead localisation

The surgical target was the STN. DBS systems from two companies were implanted: Medtronic Inc. Neurological Division, USA (octopolar directional leads, SenSight^TM^ model 33005) or Boston Scientific, USA (octopolar directional leads, Vercise^TM^ model DB-2202). Electrodes were implanted as described before^10^, connected to temporary lead extensions and externalised through the temporal or frontal scalp.

### 2.3. Stimulation and data recording

Recordings were performed ON dopaminergic medication, four to six days postoperatively while leads were externalised. Monopolar stimulation was delivered using an ISIS neurostimulator (inomed Neurocare Ltd., Germany) and referenced to a self-adhesive electrode attached to the patients’ back. Stimuli comprised symmetric, constant-current, biphasic pulses (60 µs, negative phase first). LFPs were amplified and sampled at 4096 Hz in unipolar mode using a multi-channel amplifier (TMSi Saga, TMSi International, Netherlands), with common mode rejection and custom-written software.

### 2.4. Experimental paradigm

We first performed a contact testing where we delivered 25 single pulses at 4 mA spaced 2-2.5 seconds apart to all LFP channels in sequence while recording from all other channels. The channel that elicited largest ERNA amplitudes was chosen for stimulation. We recorded LFPs during sleep and spontaneous naps.

Sleep recordings were conducted with patients 1 and 2 in the evening (between 6pm and 10pm). The two patients were laying on a recliner armchair and bed, respectively, in a dark room with minimal noise and encouraged to fall asleep naturally. Single stimulation pulses were delivered every 2-2.5 seconds with concomitant LFP, EEG (Fz, F3, Cz, C3, Pz, P3), EOG (horizontal and vertical), ECG, EMG (submental) and accelerometer (upper and lower limb) recordings.

Nap recordings from patients 3 and 4 were conducted in the afternoon (between 2pm and 5pm) when patients were comfortably seated in an armchair. Patients fell asleep spontaneously without instructions and were woken up twice (patient 3) and once (patient 4). Single stimulation pulses were delivered every 2-2.5 seconds with concomitant LFP, EEG (Cz, C3, C4, CPz, CP3, CP4), EMG (bilateral forearm flexors and extensors) and accelerometer (bilateral hands) recordings. All participants confirmed that they fell asleep during testing and this was consistent with clinical observation.

### 2.5. Signal processing

Data analysis was performed in MATLAB (version 2023b, Mathworks, Massachusetts, USA) and Python.

#### 2.5.1. Signal processing: ERNA analysis

Continuous LFP signals were high-pass filtered at 1Hz and ERNA was analysed from the contact that showed largest ERNA amplitudes^10^. We selected a period of 40ms following every DBS pulse, as this was shown to cover the duration of ERNA in previous studies. Data within 1.2 ms after the positive deflection of the DBS pulse was removed from analysis, as this was contaminated by artefact. Subsequently, we linearly detrended the selected signal, applied a second-order low-pass filter at 700Hz and upsampled the signal using spline interpolation by a factor of 10. Different features of the ERNA were extracted after each stimulation pulse: ERNA amplitude was defined as the difference between the first peak and the next trough, latency as the time between the first peak and the peak of the previous stimulation pulse, width as the half-prominence width of the first peak and duration as the time between the last peak with a minimum width of 0.5 ms and minimum prominence of 10 µV, and the peak of the previous pulse^11^ (**Figure 1A**). Spectral power was estimated from the same contact using continuous complex wavelet transform (10 wavelet cycles) for the following frequency bands: delta (1-4Hz), theta (5-7Hz), alpha (8-12Hz), sigma (13-16Hz), beta (13-30Hz) and gamma (31-40Hz). Spectral power was normalised to the average power between 1 and 40 Hz and averaged over 1-second epochs before (“pre”) and after (“post”) every stimulation pulse. **Figure 1B-E** shows traces of spectral components of the 1-second window before a given DBS pulse (“pre”).

**Figure 1.**
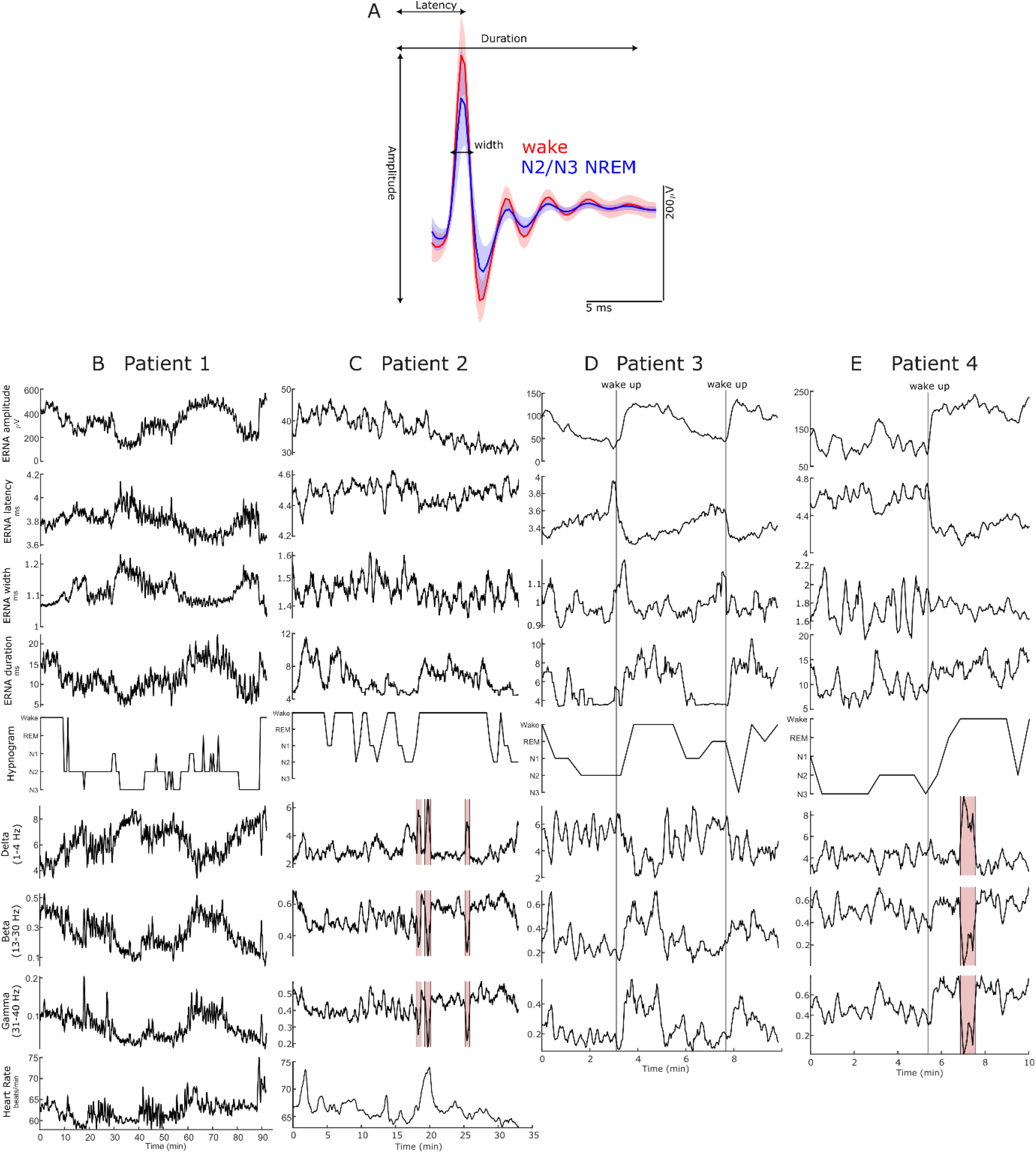
ERNA and spectral LFP power modulation during sleep. **A**. ERNA waveform during wakefulness and N2/N3 NREM sleep in patient 1 (mean ± SD). **B.-E**. ERNA features, sleep stages, spectral LFP power bands and heart rate (in B+C) are shown during sleep/nap for patients 1 to 4. Red rectangles in C and E denote movement artefacts which were excluded from the correlation analysis. Vertical lines in D and E denote the moment when patients were woken up.

#### 2.5.2. Signal processing: hypnograms

We removed stimulation artefacts in the EEG recordings and interpolated the missing signal. Subsequently the YASA toolbox was used for sleep stage labelling^12^, which outputs a classification for each 30s epoch. In patients 1 and 2, one EOG, a submental EMG and one EEG contact were used as inputs for YASA. In patients 3 and 4, one single EEG channel was used as input. To increase the accuracy of labelling, we conducted sleep stage categorisation twice with different EEG channels (Cz-Fz and Pz for patients 1 and 2; Cz and CPz for patients 3 and 4). Epochs were only considered for further analysis if the classification was consistent when different EEG channels were used, and the decoding probability of the YASA model exceeded 50% in both cases. Patient 2 was excluded from this final stage of the analysis (sleep stage classification) as this threshold was not met to reliably decode N2/N3 states based on EEG measurements and YASA.

#### 2.5.3. Signal processing: sleep stage classification

We trained support vector machine, logistic regression and random forest models using ERNA parameters and/or spectral power features for each participant to distinguish N2/N3 NREM from wakefulness. When ERNA features were considered, models were trained using parameters of single ERNA events after each stimulation pulse in patients 1, 3 and 4, as well as parameters over a moving average of ten ERNA events (10 stimulation pulses) in patient 1. Given the high degree of feature correlation, for instance between ERNA amplitude and latency, we used t-distributed stochastic neighbour embedding (t-SNE) for dimensionality reduction prior to fitting the classifier. For comparison, the average spectral power of predefined frequency bands during pre and post epochs, i.e. 2 seconds around each stimulation pulse, were also used in separate models. 5-fold cross-validation was used to evaluate the performance of each model, while the outputs from YASA are used as a gold-standard. Classification accuracy and area under the curve (AUC) of the receiver operating characteristic are reported for the best model (based on AUC) in **Figure 3**. Furthermore, hyperparameter optimisation is utilised, but only for the best feature and model combination achieved during the testing phase, to give an overall best classifier.

### 2.6. Statistics

A Shapiro-Wilk test was used to assess normality and subsequently we performed a two-sample t-test or Wilcoxon rank sum test. When correlations are reported, we performed a Spearman rank correlation and p-values are reported after FDR correction.

## 3. Results

### 3.1. ERNA is modulated by sleep onset, transition between sleep stages and awakening

We investigated ERNA and spectral changes in STN LFPs during sleep specifically to test the hypothesis that ERNA changes with sleep. As N1 is generally difficult to distinguish from wakefulness and physiologically distinct from other NREM stages, and REM sleep was detected only briefly if at all due to our relatively short recordings, we focused our analysis on N2 and N3 NREM stages vs wakefulness, as in a recent study^6^. In patient 1, we recorded a full sleep cycle (**Figure 1B**) and found that ERNA amplitude (p<.001; Cohen’s d=1.19) and duration (p<.001; d=1.11) decrease from awake to N2/N3, while latency (p=.007; d=0.56) and width (p<.001; d=1.35) increase (**Figure 2A**). Spectral features in STN LFPs demonstrated expected classical changes in canonical frequency bands with delta activity increasing from awake to N2/N3 (p<.001; d=1.03); beta (p<.001; d=1.08) and gamma (p<.001; d=0.79) activities decreasing. Accordingly, ERNA amplitude and duration were positively correlated with theta to gamma power, heart rate and sleep stages, with inverse relationships for latency and width (all p<.001).

**Figure 2.**
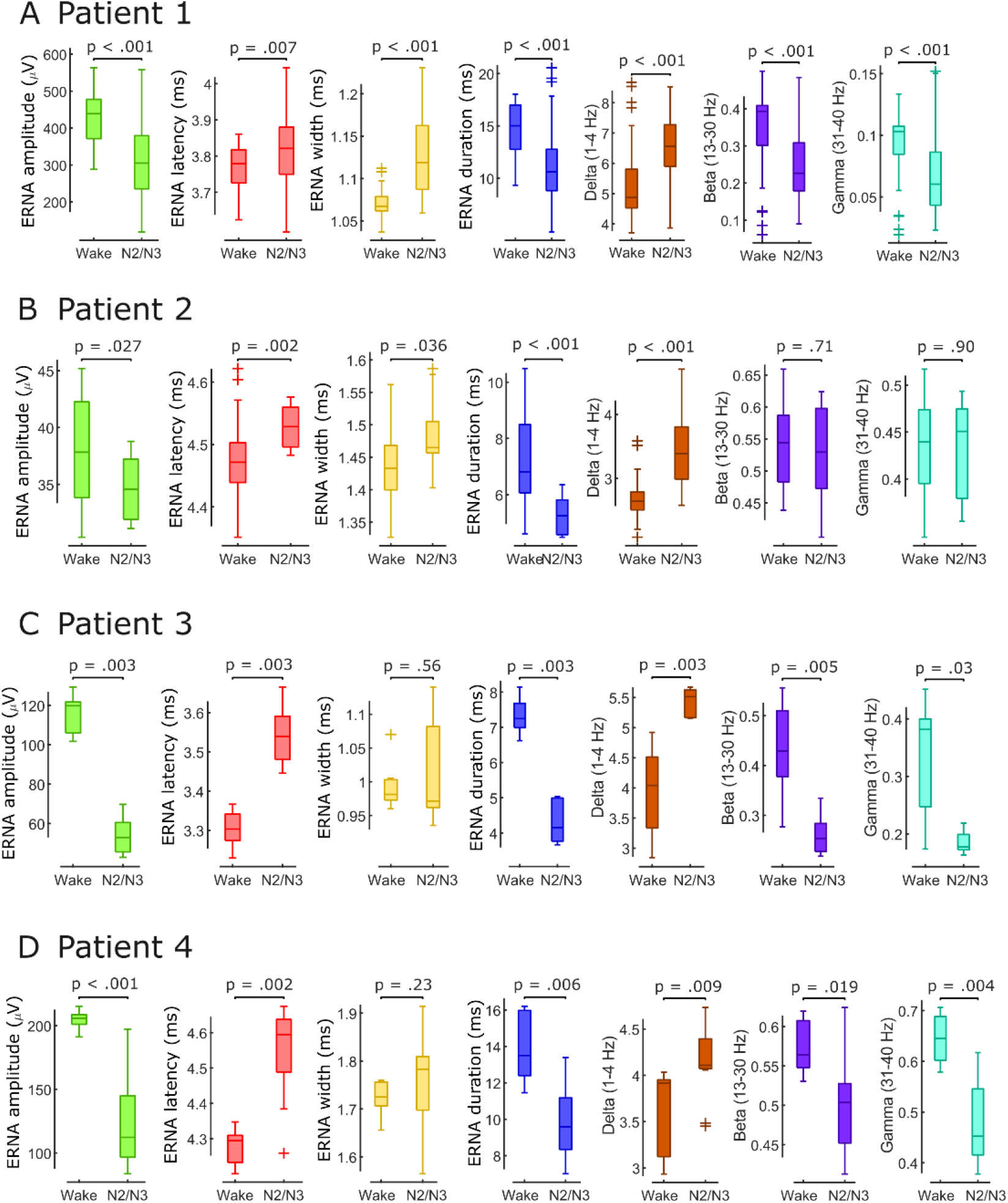
ERNA and spectral LFP features differ between wakefulness and NREM sleep. **A-D.**ERNA features and spectral LFP power bands change between wakefulness and N2/N3 NREM sleep in patients 1-4.

**Figure 3.**
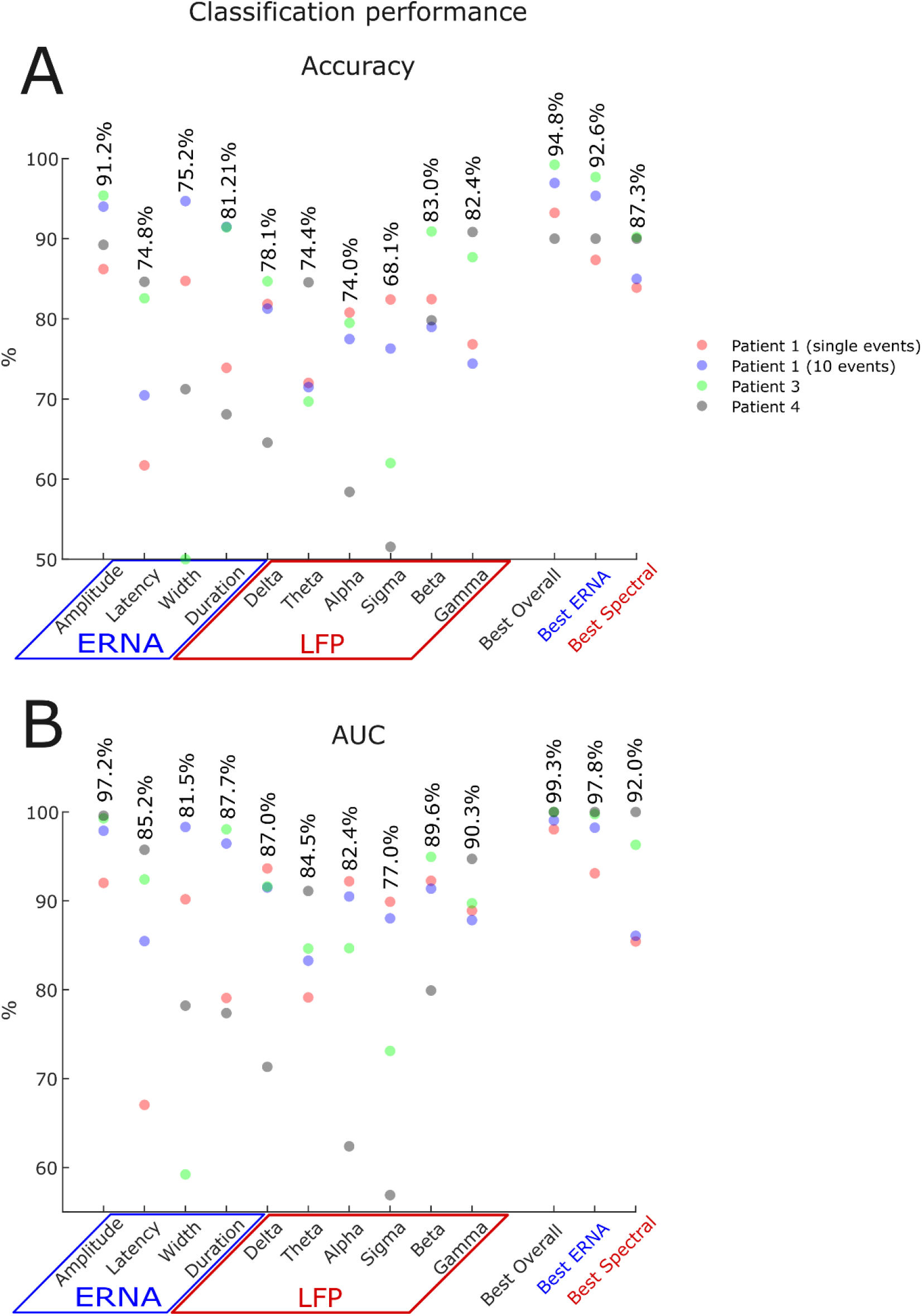
Classification performance. (A) N2/N3 NREM vs wake classification performance shown for patients 1 (using single ERNA events and 10 consecutive events which equals ∼ 25 seconds), 3 and 4. (B) Averages for the 5-fold cross validation were shown for every feature. ‘Best ERNA’ refers to the best model developed using only ERNA features, ‘Best Spectral’ refers to the best model using spectral features, and ‘Best Overall’ refers to the best model using all features (spectral and ERNA features).

In patient 2 we recorded the first ∼ 33 minutes of a sleep cycle (**Figure 1C**) and found similar ERNA changes: amplitude (p=.027; d=0.80) and duration (p<.001; d=1.32) decreased from awake to N2/N3, latency (p=.002; d=0.99) and width (p=.036; d=0.93) increased (**Figure 2B**). While delta activity increased from awake to N2/N3 (p<.001; d=2.00), classical beta (p=.71; d=0.19) and gamma (p=.90; d=0.07) power suppression was not observed in this patient manifesting as negative correlations between these frequencies and ERNA amplitude (all p<.001). This may explain why EEG-based sleep stage identification did not yield consistent labelling.

Patients 3 and 4 were recorded during spontaneous daytime naps which allowed us to record ERNA during cued awakenings (**Figure 1D+E**). In both cases, YASA did identify periods of NREM sleep despite the short nap durations. ERNA changed with the identified sleep stages: amplitude (patient 3: p=.003; d=5.98; patient 4: p<.001; d=2.70) and duration (patient 3: p=.003; d=5.31; patient 4: p=.006; d=1.97) decrease from awake to N2/N3 (**Figure 2C+D**) while latency increases (patient 3: p=.003; d=3.72; patient 4: p=.002; d=2.50) and width remained unchanged (patient 3: p=.56; d=0.36; patient 4: p=.23; d=0.33). This was coupled with corresponding spectral changes – delta power increases (patient 3: p=.003; d=2.50; patient 4: p=.009; d=1.30) and beta/gamma power decreases (patient 3: p=.005; d=2.19; p=.03; d=1.85; patient4: p=.019; d=1.30 p=.004; d=2.17) – resulting in positive relationships between ERNA amplitude, duration and alpha to gamma activity and negative relationships between ERNA amplitude, duration and delta and theta power (all p<.001). Of note, cued awakenings resulted in immediate marked changes of ERNA and spectral power supporting their modulation by the sleep/wake cycle.

### 3.2. ERNA outperforms subthalamic spectral LFP features in terms of classifying NREM vs. wakefulness

We next investigated classification performance using machine-learning (ML) models based on ERNA features, spectral frequency bands or a combination of these to distinguish N2/N3 NREM from wakefulness. We trained participant-specific models based on parameters from single ERNA events (and additionally in patient 1 based on parameters averaged for 10 consecutive events using a sliding window) and evaluated the classification accuracy and AUC (**Figure 3 and Table 1**). Across the three participants, we found that the classification accuracy and AUC (averaging at 91.2% and 97.2%, respectively) were highest for the model with ERNA amplitude as inputs, and were only exceeded by the model with feature combinations, which always included ERNA amplitude. Importantly, decoding based on ERNA amplitude is superior to all spectral components individually and in combination.

## 4. Discussion

We collected cortical and subcortical neural recordings from four participants with Parkinson’s disease and found pronounced ERNA modulation during sleep which correlated with concomitant spectral power and heart rate changes. In one participant we recorded a full sleep cycle where ERNA parameters consistently changed with respective sleep stages. Finally, we showed that ML models based on ERNA features achieved higher performance in classifying N2/3 NREM versus wakefulness than models with spectral features, with combinations of ERNA and spectral features achieving the best performance. Our findings are consistent with previous studies showing that the response of cortical activity to external magnetic stimulation is modulated by sleep^13^. This is the first study that shows STN ERNA modulation during sleep and that ERNA can be used to detect sleep stages. This can be used to develop sleep-aware adaptive DBS and improve symptom control during sleep.

### 4.1. Mechanism underlying ERNA modulation during sleep

ERNA has been suggested to result from inhibitory-excitatory reciprocal connections between the external globus pallidus (GPe) and STN^10,14^. Furthermore, ERNA was shown to align with rhythmic inhibitory synaptic input to STN from prototypic GPe neurons with ERNA amplitude being positively correlated with the potency of inhibition^15,16^. GPe neurons in turn were shown to decrease firing activity during slow wave sleep, which is consistent with the lower ERNA amplitudes during NREM sleep observed here^17^.

In addition, a recent study showed that electrical impedance in the human limbic system changes with the circadian cycle, with the limbic brain tissue impedance reaching a minimum value during NREM sleep, intermediate values in REM sleep, and rising throughout the day during wakefulness^18^. The brain tissue impedance may give rise to changes in LFPs^19^ and the volume of tissue activated by electrical brain stimulation^20^. Further research is required to investigate whether electrical impedance in the basal ganglia changes with sleep, and whether this contributes to the changes in ERNA observed here.

### 4.2. Practical use of ERNA for adaptive DBS during sleep

Beta activity is the most developed feedback marker for closed-loop DBS but beta-triggered adaptive DBS with a constant threshold may not be optimal for sleep. The overall beta reduction during sleep would lead to reduced stimulation intensity which would likely be problematic for patients who benefit from overnight stimulation and result in recurrence of PD symptoms^6^. To overcome this, the emerging field of sleep-aware adaptive DBS advocates for adjusting stimulation parameters or adaptive control algorithms with sleep stages. This relies on accurate identification of sleep stages. Most previous studies used spectral features from cortical or subcortical signals reporting accuracies ranging from 70.8% to 95.5%^4–8^. Subthalamic ERNA-based classification accuracy reported here is comparable with that of best performance previously reported, despite only using one hemisphere for decoding and without the need for additional hardware implants. Furthermore, spectral features in EEGs or LFPs are subject to movement artefacts and large cross-patient variations. As recently reported, in some patients subthalamic and pallidal beta power may even increase during sleep complicating its use for sleep stage decoding^21^. This may also explain the absence of beta/gamma suppression in patient 2 (**Figure 1C+ 2B**) while subjectively the patient confirmed they fell asleep, delta activity increased and both ERNA amplitude and heart rate decreased, suggestive of falling asleep. In comparison, ERNA comes with the advantage of high signal-to-noise ratio, being robust against movement artefacts and easy to extract from the time series as it does not rely on real-time time-frequency decomposition. A recent study showed that ML can track ERNA with ∼ 99% accuracy^22^. However, given the magnitude of the ERNA when leads are well-placed, a much simpler algorithm for extraction as we used here may suffice. ERNA-based sleep stage decoding could be used to change the threshold for beta-burst triggered adaptive DBS or to modulate stimulation intensity or frequency of continuous DBS. In the first case, single stimulation pulses could be administered between DBS bursts to evoke and measure ERNA. Importantly, beta-burst triggered adaptive DBS allows for vesicle replenishment between bursts and does not decrease ERNA amplitudes and frequencies as observed during continuous DBS^10,23^. ERNA-based DBS titration with continuous DBS may still work as ERNA amplitudes at steady state may decrease further during sleep. This needs to be explored in subsequent studies.

### 4.4. Limitations

To accurately extract ERNA, an amplifier with a sufficiently high sampling rate is required. Here, we used an amplifier with a sampling rate of 4 kHz and we previously quantified ERNA with data sampled at 2 kHz^23^. The Medtronic Percept device is already capable of 1 kHz sampling, however, not in the commercially available version. While higher sampling rates would be desirable, this may suffice to extract ERNA from well-placed leads. With the advent of rechargeable and next-generation devices, higher sampling rates that are economic may only be a matter of time.

A challenge in awake vs sleep classification is the lack of a ground truth given the interrater variability between clinicians and in part low decoding probabilities of existing algorithms such as YASA. To mitigate this effect, we only considered N2/N3 NREM epochs if YASA with two different EEG channels as input yielded consistent results with >50% decoding probability.

### 4.4. Conclusions

This is the first study to report ERNA modulation during sleep onset and sleep stages. We found high decoding accuracy to separate N2/N3 NREM from wakefulness with ERNA amplitude surpassing all spectral features which may pave the way for future personalised sleep adaptive DBS.

## Data availability

All data will be shared on the MRC BNDU Data Sharing Platform (https://data.mrc.ox.ac.uk/).

## Acknowledgements

Conceptualization: C.W., H.T., A.P. Methodology: C.W., H.T., A.P., T.G.S. Software: C.W., T.G.S., A.P., X.G. Validation: C.W., H.T., A.P., T.G.S. Formal Analysis: C.W., T.G.S. Investigation: C.W., T.G.S, S.H., F.R.P, L.W., S.Y., R.S. Resources: H.T., A.P., H.H., R.S., A.M., A.P., A.R., A.O., M.H., F.M., E.P., K.A. Data Curation: C.W., T.G.S, S.H., F.R.P, L.W., S.Y. Writing – Original Draft: C.W., H.T., T.G.S., A.P. Writing – Review and Editing: C.W., H.T., T.G.S, A.P., S.H., F.R.P., L.W., S.Y., M.H., F.M., E.P., K.A. Visualisation: C.W., T.G.S. Supervision: H.T., A.P. Project Administration: C.W., H.T., A.R., F.M., M.H., E.P., K.A. Funding Acquisition: H.T., S.H., K.A., E.P.

